# Marked point process filter for clusterless and adaptive encoding-decoding of multiunit activity

**DOI:** 10.1101/438440

**Authors:** Kensuke Arai, Daniel F. Liu, Loren M. Frank, Uri T. Eden

## Abstract

Real-time, closed-loop experiments can uncover causal relationships between specific neural activity and behavior. An important advance in realizing this is the marked point process filtering framework which utilizes the “mark” or the waveform features of unsorted spikes, to construct a relationship between these features and behavior, which we call the encoding model. This relationship is not fixed, because learning changes coding properties of individual neurons, and electrodes can physically move during the experiment, changing waveform characteristics. We introduce a sequential, Bayesian encoding model which allows incorporation of new information on the fly to adapt the model in real time. A possible application of this framework is to the decoding of the contents of hippocampal ripples in rats exploring a maze. During physical exploration, we observe the marks and positions at which they occur, to update the encoding model, which is employed to decode contents of ripples when rats stop moving, and switch back to updating the model once the rat starts moving again.

## Introduction

Neuroscience experiments are increasingly focused on the coordinated activity of neural populations and how it represents information about biological and behavioral signals. Statistical models of the population neural code allow information about these biological or behavioral signals to be inferred from the simultaneously observed spiking activity of the population. Neural population activity is typically observed extracellularly, with electrodes picking up multi-unit activity whose neural identities are unknown *a priori*. Knowing or estimating the neural identity of spikes is generally important for understanding the coding properties of individual neurons, but there has been increasing interest in statistical models of the coding properties of populations that do not require a spike estimation, or sorting, step. If the nearby neurons code for similar features [1], a simple statisical model may be created directly from unsorted multi-unit activity that can decode nearly as well as models using sorted single units [2, 1, 3, 4, 5, 6], making real-time decoding with spikes feasible. However, in cases where nearby neurons do not encode similar features, simple models based on multi-unit activity perform poorly compared to separate models for each neuron, requiring a computationally expensive spike sorting step [7], before fitting a model that collectively describe the population activity. For example, hippocampal activity recorded on a single tetrode will often yield multiple place cells whose place fields do not overlap at all, requiring spike sorting to extract spatial information from population data. New experimental designs that require closed-loop estimation and stimulation necessitates a new approach toward modeling population activity in a real-time setting.

The point process framework has been used extensively to model the activity of individual neurons [8, 9, 10, 11] once spikes have been sorted, requiring two independent steps. For population activity, modeling each spike train as separate point processes and including spiking history of other neurons in the population as covariates allows the modeling of the dependencies in the population [11, 12, 13], though cannot account for biologically relevant instantaneous joint spiking events [14, 15, 16]. An extension of the point process framework being increasingly applied to neural data analysis, the marked point process model [17], which can account for instantaneous joint spiking events [18], and has also allowed characterization of spikes according to waveform and the modeling of the populations to be done in single, integrated step. This approach, sometimes known as clusterless encoding and decoding [19, 20, 21, 22], where a joint model for the firing intensity dependent on waveform and stimulus features, behavioral variables or in hippocampal recordings, spatial position, the joint mark intensity function (JMIF) is estimated simultaneously in one step, allowing a streamlined real-time analysis pipeline.

While spike sorting methods have also improved in speed and accuracy [23], there are other advantages the clusterless method might have for modeling and decoding applications. Sorting introduces decoding errors when neural identities are determined by hard decision boundaries, and spikes with ambiguous waveforms may be classified incorrectly, in contrast to probabilistic assignment of spikes [24, 25, 19, 20]. Ambiguous waveforms in the clusterless method are automatically less informative to the decoder [20], making them less likely to cause large errors, and more suited to real-time decoding applications.

However, there are still practical issues which must be addressed with the clusterless method. First, the class of models typically used for the JMIF have been non-parameteric, resulting in an ever-increasing model size as new data becomes available. When tens to hundreds of thousands of spikes from a population are available, the computational cost of computing the model estimates may preclude real-time and closed-loop analyses. Second, neural firing properties are generally non-stationary; this may be due to changes in the spike waveforms from individual neural due to mechanical instability of electrodes or changes in background noise statistics, or due to biologically relevant changes in tuning properties of individual neurons or the population as with learning and plasticity [26].

For models of sorted spike train, a number of approaches for these issues have been addressed in part by spike sorting and clusterless algorithms. In online, real-time settings, spiking and behavioral data arrive as they become available, and non-parametric estimation methods like the kernel density estimate, linearly increase in size with additional data. Sodkomkham et al [21] have added heuristic rules for merging new spikes to existing clusters in the position-mark space based on the Mahalanobis distance between them to keep model size from growing as more spikes arrive, making decoding possible in real time even when training data is large. However, the fitting procedure treats space and waveform features equally without accounting for nonuniform sampling of space, making accommodation for nonstationarity of the JMIF more challenging. Nonstationarity of spike waveforms due to the mean waveform changing over longer timescales, possibly due to sometimes abrupt electrode drift and/or biological or background noise changes [26], can cause sometimes drastic changes in the statistics of the signal. Spike sorting algorithms account for nonstationarity of waveform features and can accommodate appearance and disappearance of neurons [27, 28, 29], but do not support online update of the model. Nonstationarity manifests also in non-spike sorted brain computer interface applications, and investigators have automated the recalibration by taking advantage of the relatively invariant low-dimensional manifold for which the neural spiking noisely codes for [5] or by taking advantage of very long histories of neural-to-kinematic mappings by using nonlinear multiplicative recursive neural networks [6]. Our contribution, the joint mark intensity function is sequentially updated mixture of Gaussians (JOYFULMoGs), builds on the marked point process framework by addressing these disparate issues using an adaptive mixture of Gaussians structure for the JMIF model, with time-varying parameters estimated under a comprehensive Bayesian framework. The mixture of Gaussians is a parsimonious choice for the model class, which requires much fewer parameters than an alternative model class such as multidimensional splines, particularly with each additional mark and spatial dimension. One application area for JOYFULMoGs is for real-time estimation of hippocampal replay events during navigation tasks. Hippocampal replay involves the reactivation of patterns of spiking activity during rest periods that occurred previously during active exploration, and the informational content of this activity likely changes throughout the course of an experiment, necessitating the need for models that can be estimated, updated and adapted in a computationally efficient manner in real time. Under JOYFULMoGs, the encoding model would be updated through each period of active exploration, and the resulting model would be used to estimate replay content during the rest periods. Note that these rest periods could be periods of sleep between experimental epochs or they could be brief pauses between exploration within an experimental epoch.

While building the full real-time decoder of hippocampal replay and deploying it in experiment is beyond the scope of this paper, we illustrate the application of JOYFULMoGs to real offline data from rat hippocampus by addressing a simpler problem to illustrate the framework. We will divide an extended period of active exploration into a sequence of shorter epochs, and alternate between adapting the encoding model in the first half of the epoch, and using that model to decode the actual movement of the rat in the second half of the epoch. The halves need not be of equal temporal duration. We call the *n*th encoding/decoding epoch I_*n*_, which contains an encoding 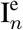, followed by a decoding 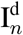, period. Both the spike waveforms (the marks) and behavior observed during 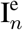 are used to update the model at the end of 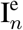, and the marks observed from during 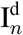 will be decoded at the end of 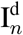 and compared with actual movements the animal made. Our model is able to handle “appearance” and “disappearance” of neurons and receptive fields, as well as drifts in receptive field location, firing rate and tuning widths and changes to the spike waveforms. The estimated posterior from 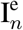 is incorporated as a prior for the same parameters in 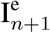. Decoding with this model is computationally inexpensive as there are only a small number of parameters, and model size is kept constant even as new data arrives. Uncertainty can be added to the posterior distribution when it is carried over as a prior in to the next 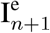, to accommodate statistical changes in receptive fields and recording quality as the experiment progresses [30, 31].

The paper is organized as follows. In the Methods section, we motivate sequential updating of the encoding model, and introduce notation used throughout the paper. We then describe the 2-step Gibbs sampling procedure, and derive the conditional posterior distributions for each of the model parameters. We also describe how uncertainty is added to the estimated posteriors when they are used as priors for the next epoch, and also how the number of clusters is updated. After describing the model, we describe a simulation of a rat running though a circular maze and simulated place field activity, before demonstrating JOYFULMoGs on dynamic simulated place fields and on actual experimental data, comparing how our method works in data settings where the data is likely to be very dynamic with changing neural representation during the experiment, and a setting where the representation is not expected to change as much.

## Methods

Real-time decoders of neural activity must incorporate continually arriving new data into the encoding model, as refitting the model from scratch every time new data arrives would be computationally expensive. Further, during the experiment, changes to the neural activity might cause some features of the encoding properties to disappear, while new features may appear, and signal and noise statistics of the spike waveforms on the electrodes may change as electrodes physically move. Fitting without regards to time would overestimate variability in the signal due to these sorts of longer-timescale trends. We propose to address these two concerns using sequential Bayesian updates to the JMIF. Since the experimental design involves interleaved periods of encoding and decoding, instead of updating with each new spike, we accumulate spikes over a period of continuous physical movement. All of the data that have arrived by the *n*th epoch, *Y*^*n*^ = {*Y*_1_, *Y*_2_, …*Y*_*n*_}, come in packets, and at the end of each 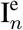, we combine the likelihood from the spikes and trajectory, *Y_n_*, observed during 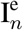, with the summary of the current state of the encoding model, the posterior from 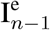, *p*(Θ|*Y*^*n*−^), interpreted as a prior, to sequentially update the encoding model, Fig. 1. The posterior distribution at the end of 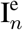 is then, by Bayes Rule,

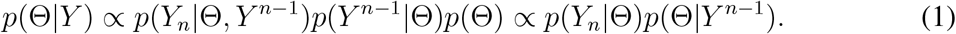

**Figure 1:**
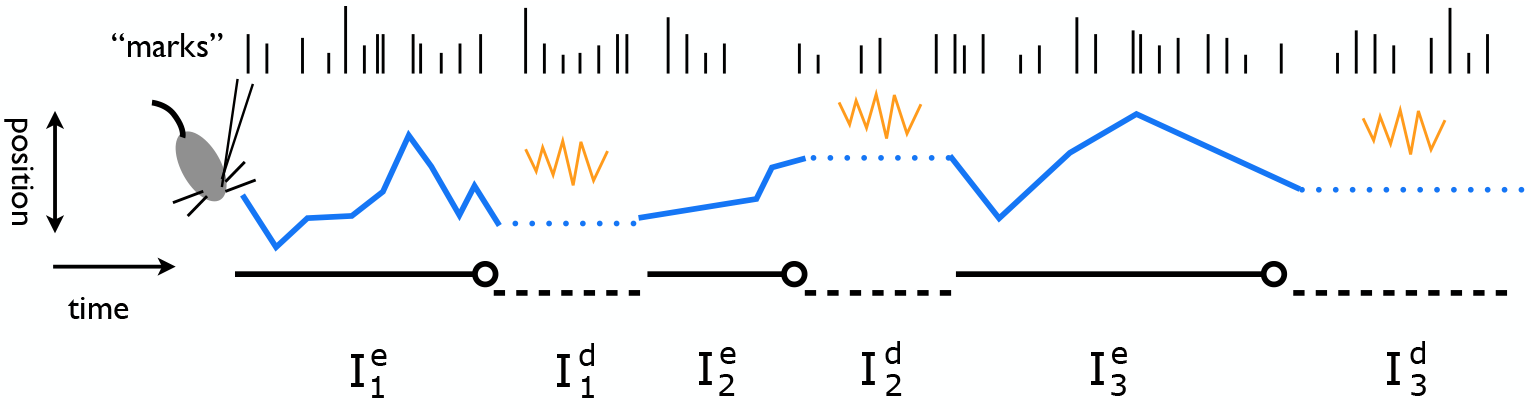
Schematic of experimental design with alternating episodes of encoding and decod-ing. Ticks at the top represent unsorted spikes or “marks”, whose varying lengths represent a waveform feature, e.g. spike amplitude. Below the marks, trajectory in blue is plotted, with solid lines representing physical movement, and dotted lines representing physical or replayed movement. Straight black lines beneath the trajectory represent time periods. Initial encod-ing model is created at the time point represented by the open circle, with prior information provided by the experimenter, using accumulated spikes and physical trajectory observed in 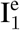, *Y*_1_. Movements or contents of hippocampal replays (sharp wave ripple as orange squiggle) occurring during 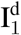 can be decoded by plugging the spikes occurring during replay into the initial encoding model. When the animal starts moving again during 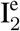, once again, spikes and physical trajectory observed during 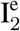, *Y*_2_, are used to the update the initial encoding model at the time point represented by the open circle, by using the initial model, *p*(Θ|*Y*^1^) as a prior to be combined with the likelihood of the data observed during 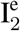. The encoding model is used again to decode movements or hippocampal replay contents occuring during 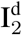.

New data is combined with the prior knowledge *p*(Θ) for the first epoch or *p*(*Y*^*n*^|Θ)*p*(Θ) for the *n*th epoch, allowing a continual update of our estimate of Θ by incorporating new data *Y* as it becomes available in real time. The posterior from 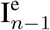 may be used as a prior for 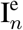 without change, or we may systematically add variance to this prior to keep the prior variance from becoming so small that new evidence provided by the most recent spikes, fails to allow the model to follow longer time-scale non-stationarities in the data.

We seek to characterize the place-specific firing of electrophysiologically recorded cells on *K* channels from rat hippocampus while it explores a maze with a trajectory *x*_*t*_. Because individual neurons with 1 or more distinct place fields presumably generated the neural activity, position and *K*-dimensional mark of spikes often appear clustered when plotted simultaneously, Fig. 2. Previous work used a non-parametric kernel density estimation to estimate this *λ*(*t*, **m**) [19, 20]. Here, we approximate it as a mixture of *C* Gaussians 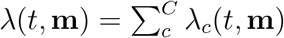, with Gaussian components

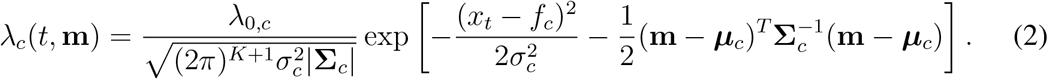

**Figure 2:**
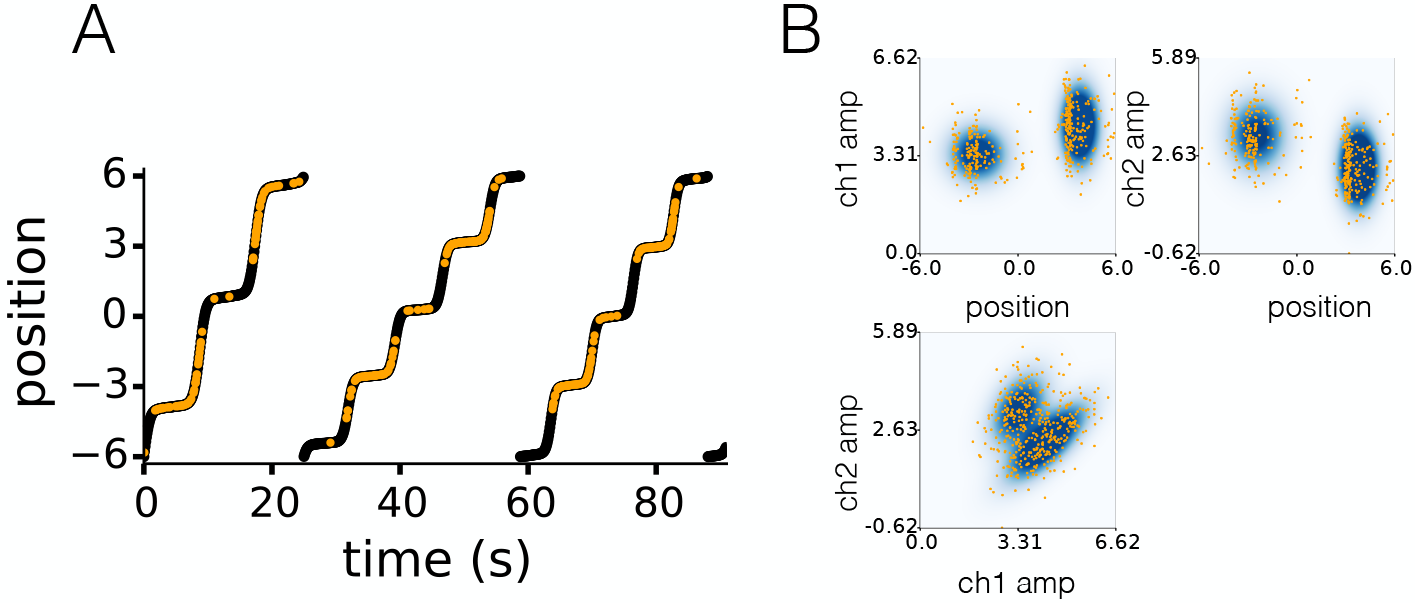
A) The trajectory of a simulated rat over a circular track, *x* parameterized from [–6: 6]. Orange dots overlayed are the time and location where an unsorted spike was observed. Rat traverses track in stops and starts. B) Top 2 rows, position vs. channel 1 or 2 and bottom row, channel 1 vs. channel 2 plot of observed marks plotted on top of marginalized JMIF. Note that the center of the scattered marks does not coincide with the fitted clusters due to the non-uniformity of the trajectory.

The hyperparameters for the centers are 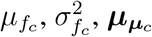, and 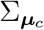, for the widths are 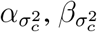,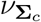 and 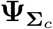, and those for the intensities are 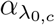 and 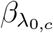

### Fitting procedure: Gibbs sampling

In 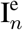, we observe *Y*_*n*_ = {Δ**N**, **M**, **x**} in *T* discrete time bins. Δ**N** is a vector of size *T* indicating how many spikes ≥ 0, are observed in each of the *T* time bins. **M** is a vector of size *T*, whose *t*th entry is a matrix of size Δ*N*_*t*_ × *K* containing the Δ*N*_*t*_ ≥ 0 marks of dimension *K* of the observed spikes. **x** is a vector of size *T*, and is the trajectory of the animal during this epoch. Initially when *n* = 0, a guess is made as to the number *M*_*G*_ and locations of (spatial-mark) clusters that are present in the data, to initialize the Gibbs sampler. We initialize with *M* components slightly larger than *M*_*G*_, and for the clusters with indices > *M*_*G*_, we randomly set their centers and widths to be random values. For the other components *c*, we use sample statistics for Θ_*c*_ as starting parameter values. However, we leave the priors for all the Θ broad to express our uncertainty in their values.

The Gibbs sampling involves repeating two steps: the stochastic allocation of all spikes into clusters given current parameter values, followed by the conditionally independent sampling of the parameters of each cluster given the assignments in the previous step [32]. Typically in Gibbs sampling in a model with parameters Θ and data *Y*, the *θ* ∈ Θ are sampled sequentially in each Gibbs iteration, from the conditional posterior *p*(*θ*|*Y*, Θ_\*θ*_). The conditional posterior is obtained by combining the likelihood *p*(*Y*|*θ*, Θ_\*θ*_) with all Θ_\*θ*_ regarded as constant, and a prior conjugate to this likelihood. After a large number of iterations (we choose 15,000) to sample an empirical posterior distribution, model parameter values are commonly read off using the empirical mean, mode or median. Here we choose the empirical median. JOYFULMoGs then uses the empirical posterior distribution as a prior for Θ when updating the model with new data in the next epoch. In order to do this, we fit the marginal empirical distributions of each parameter *θ* with the distribution conjugate to the likelihood with all Θ_\*θ*_ constant, using maximum likelihood to find its hyperparameters. This approximation of the posterior distribution by marginalization is reasonable, because when samples of a parameter pair (*x*, *y*) are examined from the marginalized empirical distribution *p*(*x*, *y*) = ∫*p*(Θ)*d*Θ_\(*x*,*y*)_), the samples often appear to have little correlation, suggesting that the joint distribution of the parameters Θ are nearly independent and the product of the marginal posteriors should be a good approximation to the joint posterior distribution^1^. We also note that not all of the parameters used in mixture of Gaussians JMIF have a likelihood with such a conjugate prior (notably *f*_*c*_ and 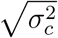). In those cases, we explain our choice of prior in the following section.

### Stochastic allocation into clusters

Conditioned on fixed parameter values Θ, a stochastic assignment of 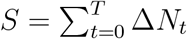 observed marks into *C* clusters is performed. Because more than 1 spike may be observed in a time bin, we introduce notation #(*t*, *w*), which maps the time bin number and the spike index within that bin to a single integer index ∈ [0, *S*]. A single row of the indicator matrix **Z** has 0s in *C* − 1 columns, and a 1 in the column corresponding to the cluster to which this mark belongs, so 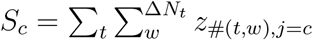 is the number of marks assigned to cluster *c*. The probability of assignment of the *w*th mark in time bin *t* to the 1st cluster is

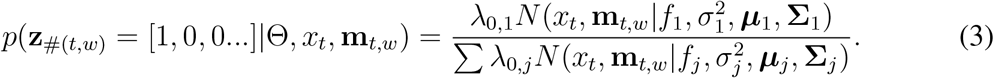

Once **z** is assigned, fitting the parameters of each individual cluster can proceed independently of all other clusters.

### Sampling the parameters Θ

Conditioned on the assignments **Z** made in the previous step, parameters Θ are sampled. The likelihood of observing Δ*N*_*t*_ marks at time *t* is

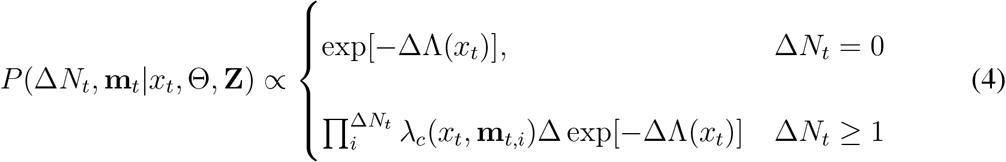

where

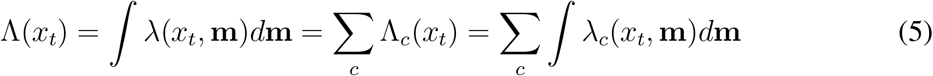

is the marginalized JMIF. We wish to estimate the parameters 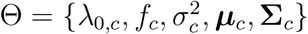. The likelihood of observing the marks {Δ**N**, **M**} given the trajectory *x*_*t*_ and cluster assignments **Z** is

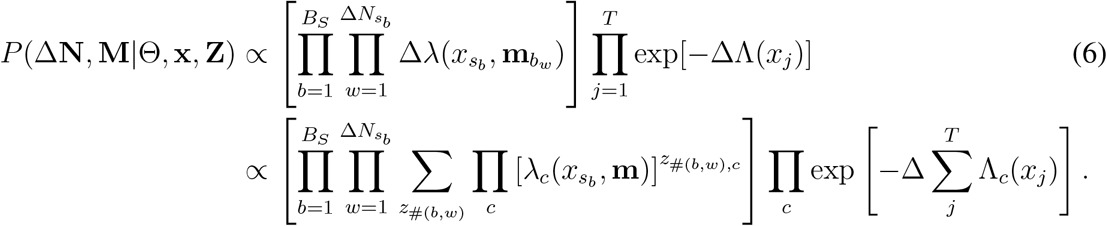

For the last exponential term, we exchange order of the product of terms evaluated at multiple timepoints (and positions) with the sum over all the Gaussian components, to obtain the last line. Rewritten, we see each component in the product can be approximated as 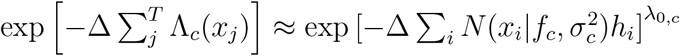, whose argument is a roughly a convolution of a Gaussian with mean *f*_*c*_ and variance 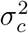 with the spatial occupation density. Convolutions are expensive to calculate at every Gibbs iteration, but this simple rearrangement allows us to approximate this term by calculating the convolution over a 2-dimensional grid of means and variances once before the start of Gibbs sampling, and doing lookups for specific values of *f* and *σ*^2^. To emphasize our computational approach, we write this term as 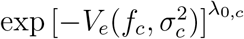, with the *e* emphasizing that, while this function changes each epoch, its evaluation is generic across all *C* clusters once the path traversed in the epoch is given. This term takes into account the nonuniform animal trajectory, Fig. 2. Absent this term, if the animal moved very slowly through a region on the outskirts of a place field of a place cell, but quickly moved through the center of a place field of the same cell, the estimate of the place field center would be biased to the outskirts. The **z**_#(*b*,*w*)_ in the exponent in Eq. 6 groups terms by cluster assignment of the spikes, so that the expressions for the *C* conditional likelihoods for each cluster only involve spikes assigned to those clusters. Thus once an assignment is made, the sampling for each of the *C* cluster parameters can proceed independently.

Samples for a parameter *θ* are drawn from the conditional posterior distribution for *θ*, which can be calculated by multiplying by the likelihood and the prior that are both calculated at fixed values of all parameters except *θ*. In what follows, we will simply write *p*(*Y*_*p*_|*θ*, **x**) as a shorthand for the likelihood in which all other parameters except *θ*, are fixed, and will assume the joint posterior to be multiplicatively separable. The sequential updating scheme depends on using conjugate priors in order that the posterior will have the same form as the prior, Fig. 3, so we parametrically characterize the marginal posteriors of each parameter using the distribution class of its conjugate prior, which in practice is often a good assumption. Not all parameters have likelihoods with a conjugate prior due to the presence of the exponential term, and for these, we will use the conjugate prior of the other term, under the assumption that the exponential term does not alter the shape of the distribution very much.

**Figure 3:**
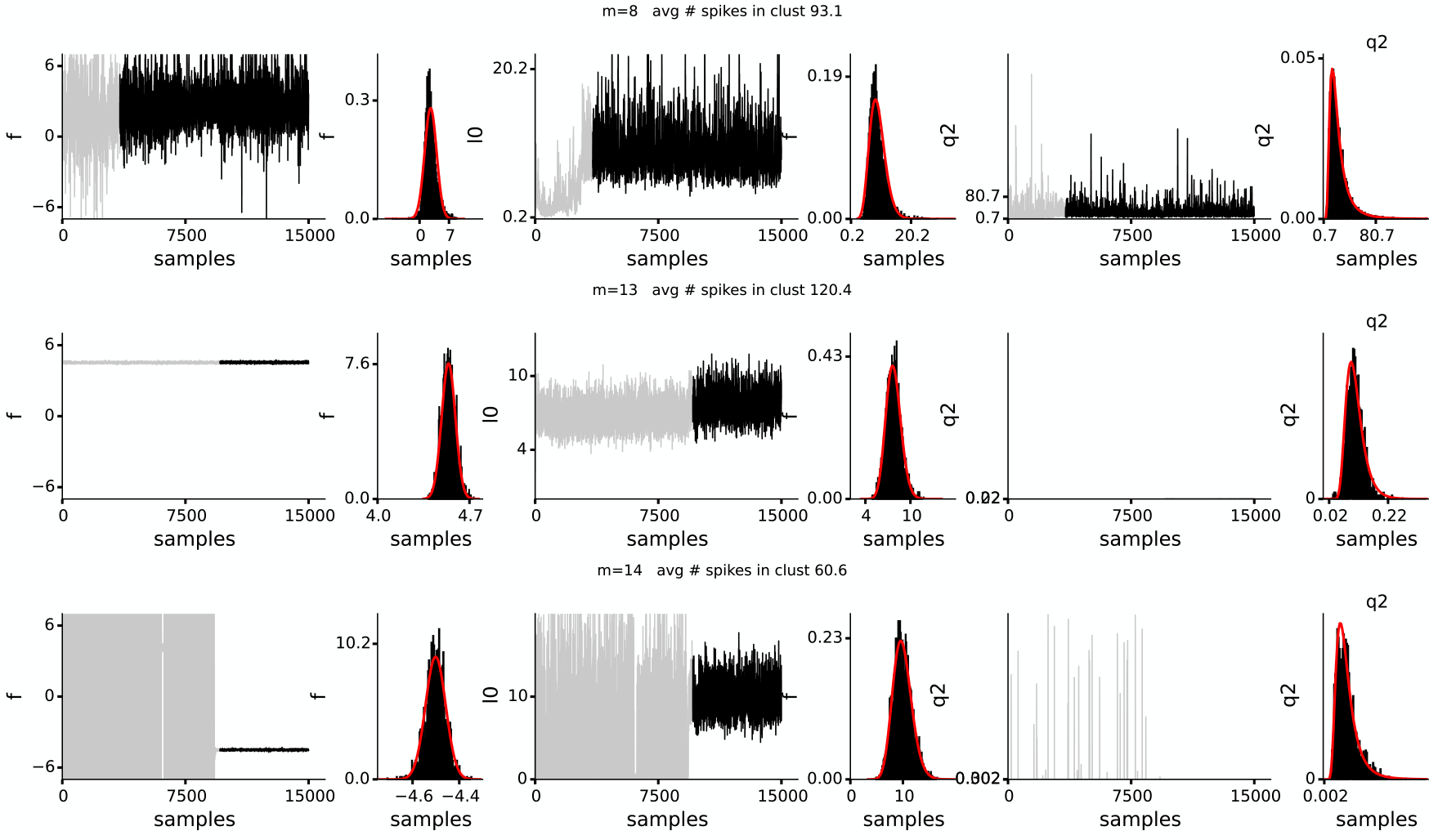
3 selected chains of spatial and intensity parameters (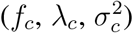)from 3 typical clusters. The parameters are shown as samples from the chain from the start of the Gibbs procedure (grey), with samples deemed to be drawn from stationary distribution overlayed in black, starting at iteration *S*_*c*_. Histograms of the stationary samples are shown to the right, with a fit by the model class of the conjugate prior (red). The assessment for stationary samples is done on a cluster by cluster basis.

### Sampling cluster intensity λ_0,*c*_

Treating all other parameters besides *λ*_0,*c*_ as constants, the likelihood Eq. 6 is 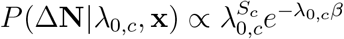, where 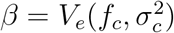. The conjugate prior is the Gamma distribution Gamma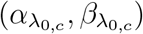, where 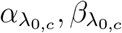 are the posterior hyperparameters from the previous epoch. The conditional posterior from which we sample is then the Gamma distribution Gamma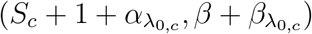, and the resulting posterior samples of *λ*_0,*c*_, are well fit by a Gamma distribution, Fig. 3, middle column.

### Sampling cluster spatial center *f*_*c*_

The likelihood Eq. 6 with terms involving *f*_*c*_ is

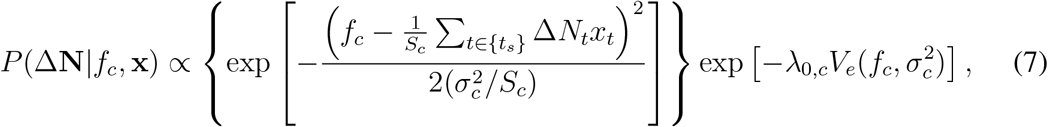

where {*t*_*s*_} = {*t*|Δ*N*_*t*_ ≥ 1}

There is no conjugate prior. However, we find that the sampled marginal posterior often is very close to a Gaussian, left column Fig. 3, which would become the prior to combine with the likelihood, so we use a Gaussian prior 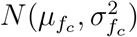 here, and obtain for the conditional posterior

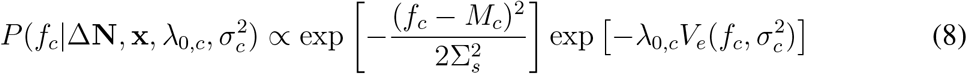

where

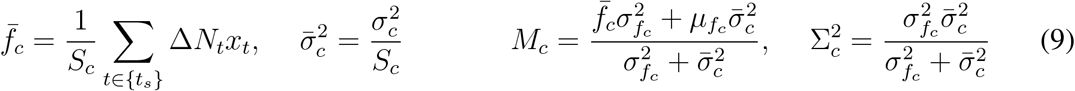

Eq. 8 is not of a known distributional class, but *f*_*c*_ can be sampled from the numerically computed cumulative distribution function, which can be done efficiently by variable-resolution sampling at selected values of *f*_*c*_, Fig. 4, and the samples fitted with a Gaussian. The variable-resolution sampling assumes the probability mass is concentrated, but not necessarily unimodal.

**Figure 4:**
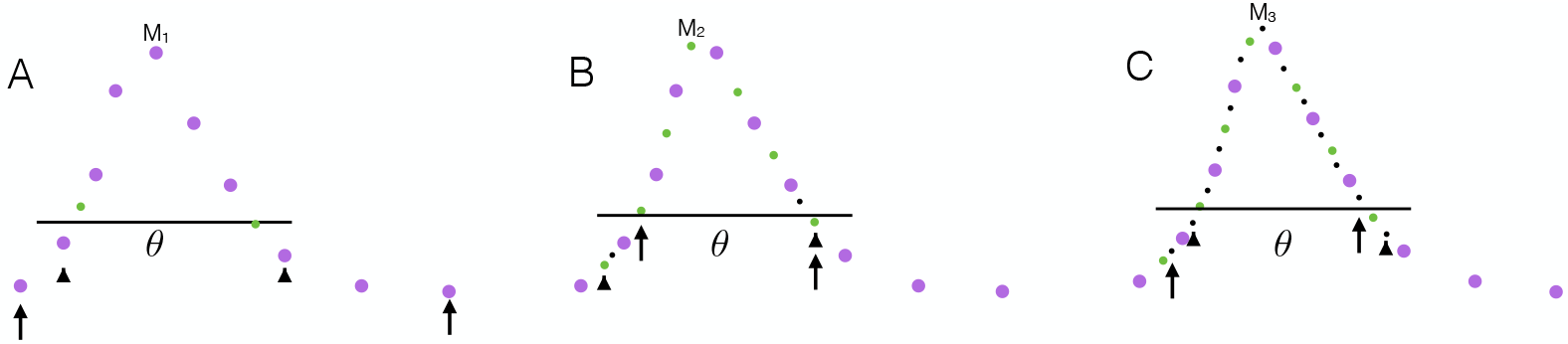
Variable-resolution sampling of conditional posterior. We first create a grid on which parameter *θ* is defined, and choose an integer *L* > 0, which determines how coarse the initial sampling on the first pass is done. A) As a first pass, the conditional posterior is coarsely sampled at every 2^*L*^th grid point, purple dots, over the range of the parameter *θ*, with *i*′ = arg max_*i*_(*θ*_*i*_|*Y*) and *M*_1_ = *p*(*θ*_*i*′_|*Y*). Starting at *i*′ and for *r* ≪ 1, search for a contiguous region of sampled values for *i*s such that *p*(*θ*_*i*_|*Y*) > *r*M_1_ (black triangles). This is the region where sampling at finer resolution should take place. B) Starting from left boundary of region found, shift by 2^*L*−1^ grid points to the left (red triangle), and sample at every 2^*L*^th grid points, until we hit the red triangle on the right. We once again find new *i*′ = arg max*i* p(*θ*_*i*_*Y*) and new limits. We repeat this process until we have a satisfactory number of sampled points within the high probability region. Now a sample can be drawn from the conditional posterior as the CDF can be easily calculated.

### Sampling cluster spatial width 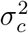

The conditional likelihood is

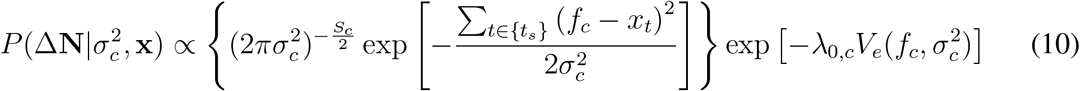

There is no conjugate prior. However, we find that the sampled marginal posterior often is very close to a inverse Gamma distribution, right column Fig. 3, so we use an inverse Gamma prior IG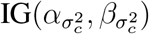 here, and obtain for the conditional posterior

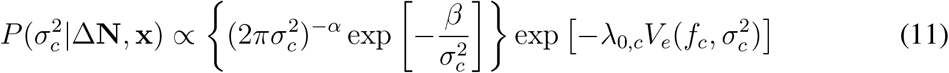

where

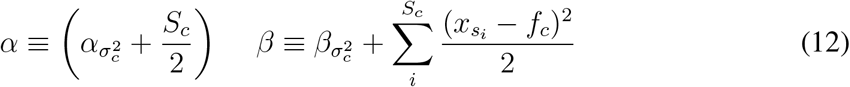

Once again, Eq. 11 is not of known distributional class, but *σ*_*c*_ can be sampled using variable-resolution sampling as used for *f*_*c*_. Here, the grid on which *σ*^2^ is defined should itself be non-uniformly spaced, but rather scaled to be wider for larger values of *σ*^2^, and variable-resolution applied to this non-uniformly spaced grid.

### Sampling cluster mark center of *µ*_*c*_

The conditional likelihood is

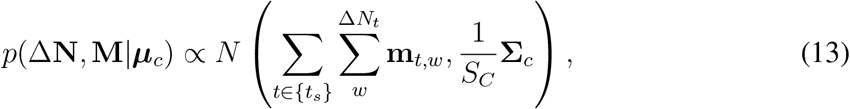

and the conjugate prior is 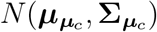. The conditional posterior is

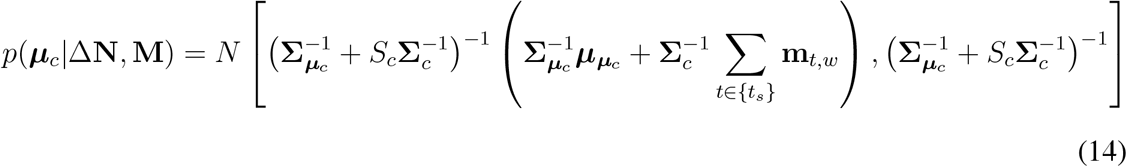

We approximate the posterior to be a multivariate normal, and estimate the hyperparameters using the sample mean and covariance of the samples.

### Sampling cluster mark width Σ_*c*_

Rewriting the Gaussian in Eq. 13 using the cyclic permutation of the trace highlights the form conditional likelihood

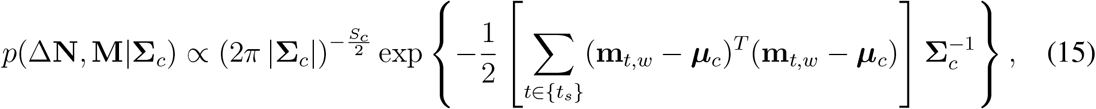

has the form of an inverse Wishart distribution, and the conjugate prior is IW(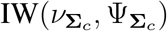). The conditional posterior is

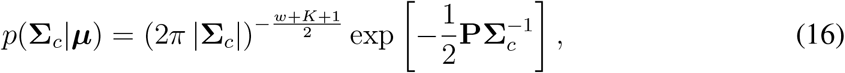

where

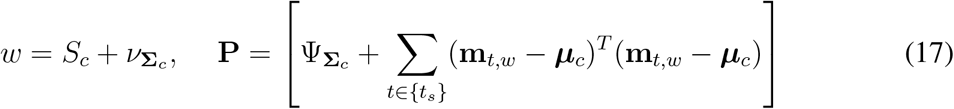

Estimating the hyperparameters of an inverse Wishart distribution using maximum likelihood is difficult, so just for **Σ**_*c*_, its hyperparameters will be the means of the hyperparameters used in the conditional posterior, Eq. 17.

### Updating number of clusters

Place fields can rapidly and dramatically change over the course of an experiment. Our approach here is not to infer the statistically optimal number of clusters to use at any point in the experiment, but ensure there is a high likelihood of observing the data, even at the expense of more than the optimal number of clusters. However, we would still like to be as concise as possible, so we use several heuristic criteria to assess whether to keep a cluster or not for the next epoch. The 1st criteria is that, on average, at least 1 data vector is assigned to the cluster throughout the Gibbs iterations. The 2nd is that the posterior distribution of the cluster center is reasonably well concentrated relative to the width of the cluster. We also require that at least 3000 iterations prior to the end of sampling be sufficiently stationary, to have some confidence that this cluster represents a well-isolated local optimum.

### Updates of the encoding model

We suggested that the joint posterior to be approximately independent in its parameters. Because of this, we implement the sequential updates by using the marginal empirical posterior directly for each parameter as the prior for that parameter in the next epoch. However, if the signal is not stationary, but its statistical properties change relatively slowly in time, this will lead to an over-estimation of the variability. Absent predictions of how the place field and mark properties may change, this is equivalent to increasing the uncertainty of the posterior when we carry it over as a prior. For example, if Θ represented a place field center, and we believe that it could change by up to 10cm in 1 minute, no matter how certain we became of Θ at the *n*th time step, we could set a minimum width for *p*(Θ|*Y*^*n*−1^) that reflects a minimum of 10cm of uncertainty added every minute. For convenience, we denote the set of parameters that describe the *c*th mixture component by Θ_*c*_. If the encoding step of two consecutive epochs are separated by time epoch *T*_*e*_, at the end of each epoch, we modify the hyperparameters as follows:

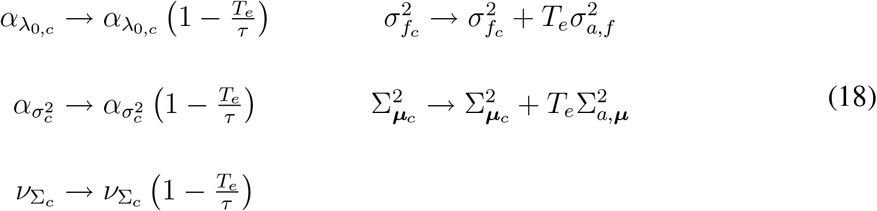

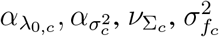 and 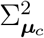 control the width of their respective prior distributions. *τ* controls the timescale in which adaptation occurs, and the 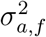 and Σ_*a*,***µ***_ the amount of spread in the uncertainty of the width of the place field and marks. For the normal distributions, the uncertainty and the centers can be adjusted independently, but for the other parameters, modifying the above hyperparameters must be done in pairs, so that the mean or mode remains unchanged.

### Simulated place fields and task

Simulations of hippocampal activity of a rat moving at variable speeds on a circular track were performed. Several neurons were defined, and their respective wave form features (marks) were generated from multivariate normal distributions whose parameters are specific to each neuron. Each neuron was then assigned one or more Gaussian place fields with accompanying intensity. Epochs defining when new data is used to update the model were either uniformly or randomly generated. We also defined low-amplitude spikes with broad tuning, and a cut-off threshold to mimic neural data used for clusterless decoding [19, 33, 20].

### Hippocampal recordings and task

Extracellular data from 12 tetrodes in CA1 and CA2 of rat hippocampus was recorded while the rat traversed a W-shaped maze. Spikes were detected at a lower threshold than for preprocessing for traditional spike sorting [7, 34], as our goal is not necessarily clean isolation of individual units, but extraction of as much information as possible from the data [35], and all above-threshold spikes were used for decoding, even if they may be a mixture of noise and spiking from sources far from the tetrode [19, 33, 20]. However, lowering the threshold at which spikes are detected leads to rapid increase in the number of spikes. This has several undesirable consequences. The stochastic allocation into clusters requires calculating the Mahalanobis distance from every data point to every cluster, so the number of computations required grows like *M* × *N*, linearly with *N*, the number of spikes identified and used for fitting. With the larger *N*, deviations of the cluster shape from Gaussian become easier to detect, and our fitting procedure may assign multiple Gaussian components to the hash cluster. Because these “spikes” may not even be of neuronal origin, investing too much time capturing the shapes of low-amplitude clusters may provide very little additional information about the physical position. However, past work suggests that these low-amplitude clusters are somewhat informative [20], so removing them entirely is undersirable, and requires us to set an arbitrary threshold for the hash, which might be difficult to do in a real-time setting. As such, we may also treat the threshold detailed in [34] as stochastic, and allow randomly allow some spikes lower than the threshold be allowed into our data. There tends to be more lower-amplitude spikes than high amplitude spikes, and this thinning allows us to keep some of these which might contain some information, without overwhelming the analysis with too many spikes that do not contribute as much information.

Rats moved through the maze in bouts of movement, interspersed with rest. The content of the hippocampal activity during movement is markedly different than during rest, with awake hippocampal replay events present in the latter [36, 37]. Because the spiking during rest may reflect not only reflect the current physical location, but internal states not accessible through monitoring of behavior, we built an encoding model using hippocampal activity and spatial location only when the animal was moving. Concatenating the periods of movement, we artificially constructed neural data of a rat in perpetual motion to test our decoder. epochs defining when new data is used to update the model were randomly generated.

The W-shaped maze is a 2-dimensional space, and the trajectory of the animal was linearized according to a scheme suggested by Deng (unpublished), so that a path going from home well H to the choice point (C), to the left (right) arm L (R) and back through C to home well H, was mapped to numerical position 0 to 1(-1) to 3(-3) to 5(-5) to 6(-6)(1, 5) and (2, 4). This mapping assigns different numbers to the same location depending on direction of movement, and requires some future information when mapping movement of animal as it leaves H, because whether the coordinates are negative or positive requires knowledge of whether it will eventually turn left or right.

### Assessment of the encoding models

We assess JOYFULMoGs by using them to decode subsequent data. We decode neural activity using the marked point process filtering algorithm described in Deng *et al* [20], and assess the decode utilizing 2 metrics, the root mean squared error, calculated by taking the mean of the squared difference of between the *maximum a posterior* of the inferred filter position and the actual trajectory, and the time under 95% posterior, *t*_95_, which is the time the actual trajectory is within the highest 95% posterior density regions.

## Results

### Sequentially accumulating evidence

JOYFULMoGs needs to solve 2 practical issues in real-time encoding and decoding. The first is the sequential estimation of a model as new data arrives, and the second is to adapt the model in cases where the underlying structure in the data changes throughout the experiment. We assess JOYFULMoGs for these 2 criteria using simulations of a rat with a single tetrode implanted in the hippocampus navigating a circular track.

In the first set of simulations, the generative models for the place fields are static, and 12 epochs all of equal duration. 8 realizations for a recording scenario with 7 neurons being recorded are generated, Fig. 5. Encoding models for each realization are built in 2 ways: first where model parameters are estimated using only data from the most recent epoch (DSCRD), discarding all past information about the model, and second where sequential updating is performed (SQUP), and the past information about the model is carried over as a prior as described in Methods. The decode performance is compared for both approaches, and also with the performance of a model using ground truth parameters (GT) the data was generated with. The DSCRD strategy produces relatively constant decode errors over successive epochs, while the SQUP strategy approaches the performance of the GT after several epochs, and outperforms DSCRD, which is not surprising as it benefits from effectively a larger amount of training data, Fig. 5. Fig. 5A-C) right, show that for stationary data, whether a model is built using all the data at once or sequentially, the resulting model is essentially the same, as expected.

**Figure 5:**
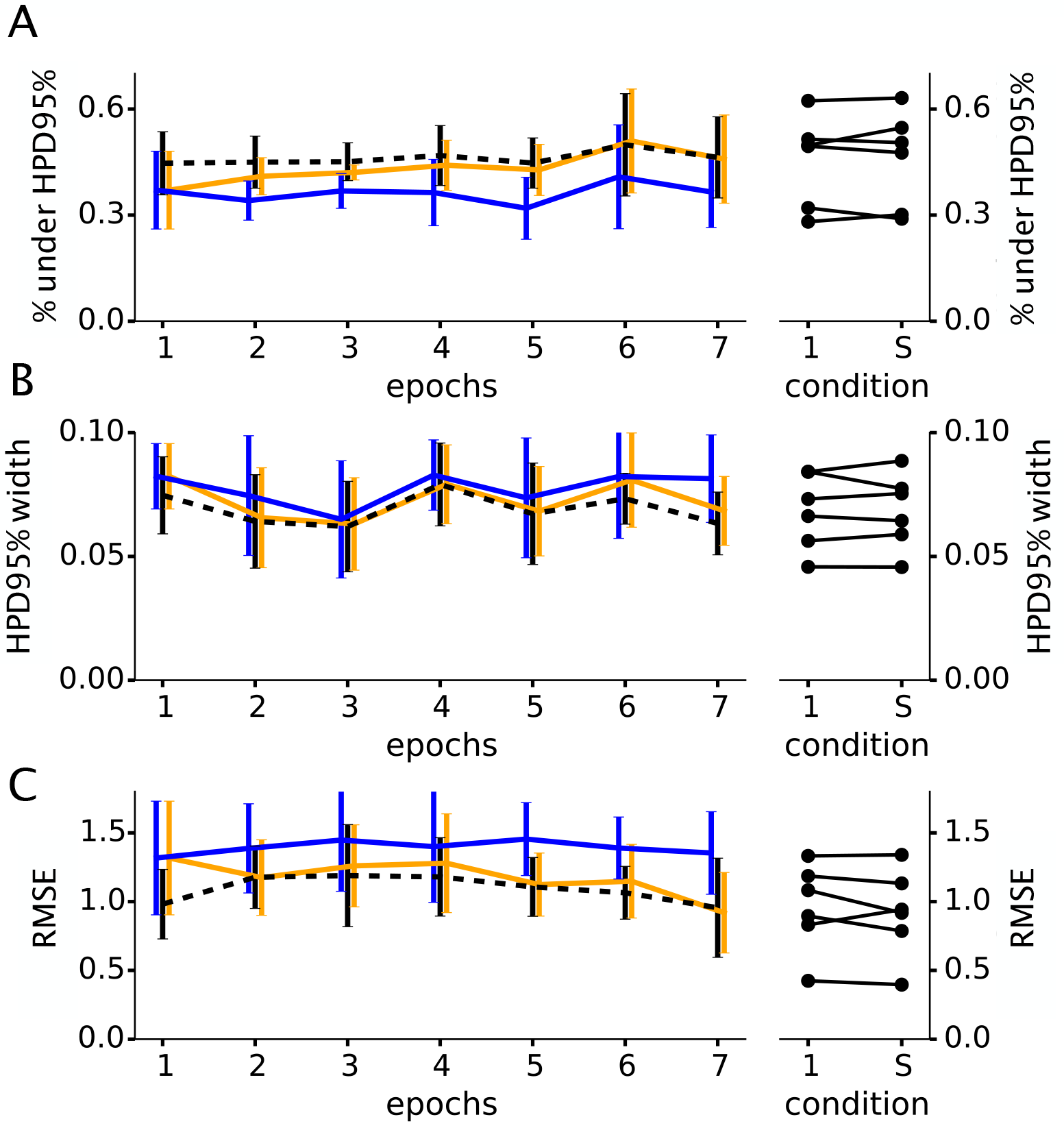
Assessment of sequentially-built encoding model on simulated, stationary data. For A-C), summary of performance metric of decode from 10 simulated experiments are plotted. Metrics are A) fraction of time actual path is under the 95% highest posterior density (HPD) area, B) the width of the 95% HPD area and C) the root mean-squared error (RMSE). Black, orange and blue lines show mean and standard deviation of decode metric for each epoch. Colors correspond to encoding model GT (ground truth parameter values at each epoch), SQUP (models built and updated sequentially) and DSCRD (no update, models built by discarding data more than 1 epoch old). Right panels show a comparison of mean decode performance of epoch 7 using model where data from all previous 6 epochs were used at once (condition 1), and the corresponding performance when model at epoch 7 was obtained by SQUP (condition S).

We note that the decode accuracy is a combination of how well our encoding model describes the JMIF, as well as how completely the place fields tile the space and how well the movement model describes actual movement. We have purposely simulated the rat’s movement to alternate between randomly occurring bouts of slow and fast walking to create regions of high spiking that are due to high occupation (slow walking) rather than place field firing, to check that our method still models the place fields correctly. Our movement model however only captures the average movement speed, and is not well matched to the alternating speeds, leading to relatively large decoding errors.

### Tracking changes in neural representation

Both biological changes, such as rapid changes in the neural representation [38, 39] and experimental changes, such as tetrode drift, may occur during the experiment. In the second simulation, we simulate place field and electrode drift, as well as more rapid changes over short timescales, such as new place fields appearing or disappearing over the course of the simulation. We will be comparing the SQUP to using a stale SQUP from several epochs previous. The stale encoding models should decode poorly near epochs where the neural representation is rapidly changing.

Fig. 6B shows the SQUP tracking the changing structure of the JMIF, as new place fields shift, appear or disappear, subjectively matching the changing features as compared to GT. Fig. 7A shows that SQUP decoder is able to keep up with rapid changes in the neural representation, and shows little difference in decode compared to the GT decoder, while the 7 epoch-old SQUP decoder shows rapid deterioration in decode quality as soon as there is rapid change in neural activity around epoch 10, Fig. 7B. We also see a gradual improvement in decode performance towards the later epochs, Fig. 7B and C, in agreement with Fig. 5, indicating that our method is able to accumulate evidence by sequential updates, while also keeping track of rapid changes present in the data.

**Figure 6:**
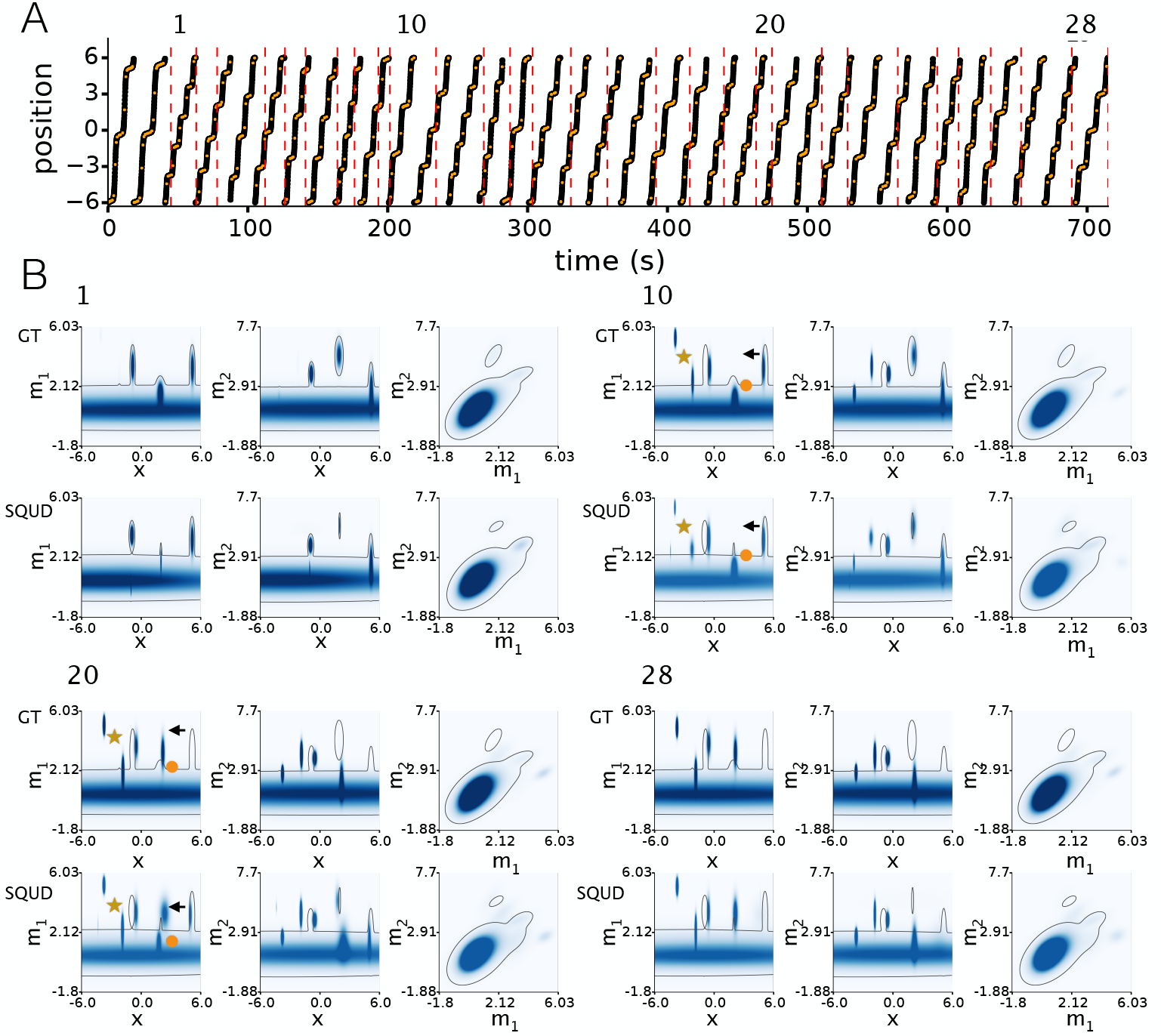
Encoding of simulated data with changing place fields. A) rat trajectory (black), overlayed with time and location of spikes (orange), with randomly chosen epochs shown by vertical, red dashed lines. The epoch numbers (1, 10, 20, 28) above trajectory correspond to epochs where JMIF is displayed in B). For each of those 4 sampled epochs, B) shows 2 rows of JMIF visualized by plotting 1 of 4 mark components against the position (left and middle), and against each other (right), for the GT (top) and SQUP (bottom), are shown. The widely-tuned component at the bottom represents hash spikes, which are low-amplitude spikes slightly above the discard threshold. The contour line shows the contour at 5% the maximum mark intensity of the first epoch, for comparison at the later epoch. 2 new place fields appear (near star), 1 place field disappears (near circle), and 1 place field rapidly shits (near arrow), around epoch 10.

**Figure 7:**
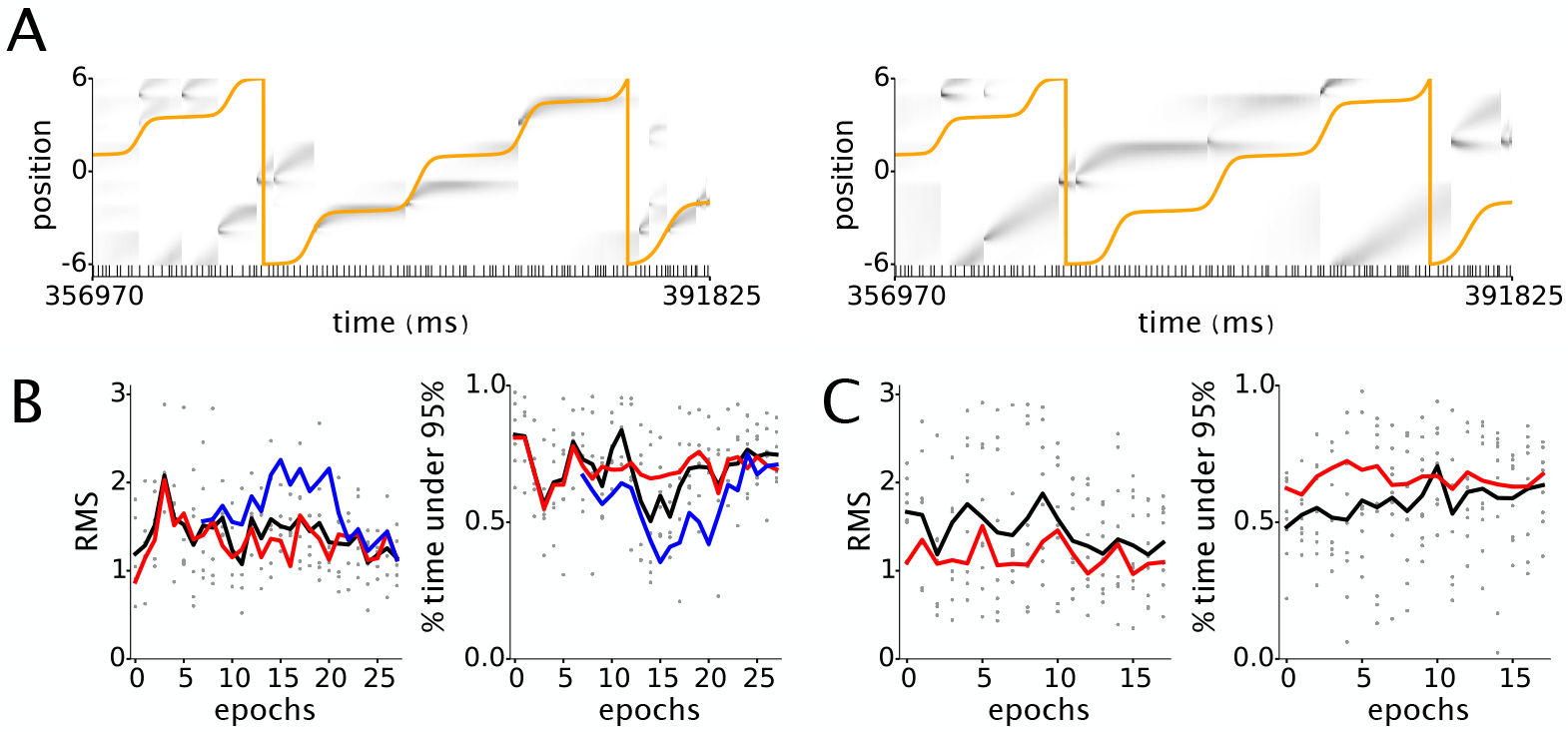
Decoding of simulated data with changing place fields. A) Decode of 1 epoch of sample realization, physical position in orange, posterior probability in grey. Decoding using SQUP (left), and stale SQUP (use model 7 epochs old). B) Summary of decodes of 9 realizations of simulated data with similar time course of changes as Fig. 6, with abrupt appearances and disappearances of place fields around epoch 10. Scatter plot (grey dots) of the RMS and % time under 95% HPD region for decoded trajectory using SQUP for each realization, while the black line is the average, the blue line the average using stale, 7 epoch-old SQUP, and red line is the average obtained using GT. C) Similar figure for another set of 9 realizations of a dataset where changes are more gradual, showing how the SQUP decode performance approaches decode performance using GT after many epochs.

### Decoding physical movements

We sequentially decode the movement of a rat in the W-track on 2 different days, one an earlier and the other a later exposure to the track, Fig. 8. The trajectory through the track is visibly different for these 2 days, with the early exposure being more variable, and sometimes deviating greatly from the sequence the arms that must be visited for a reward. For example, in Fig. 8B on the left, a correct sequence should see the animal moving through the landmarks as H-C-L-C-H-C-R-C-H-C-L-C-H…, but this is violated around times *t* = 150, 195, 310, 320, 390. Fig. 8B on the right shows the sequence being visited correctly. A cursory examination of the mark-space raster plots for the early exposure suggest that on several tetrodes, neural representation appears to be changing, while for the late exposure, noticeable changes in the neural representation are not obvious looking at the mark-space raster plots. We show that JOYOUSMoGs indeed captures the changes in neural representation when present, and sequential updates to the model leads to significant improvements in decoding.

**Figure 8:**
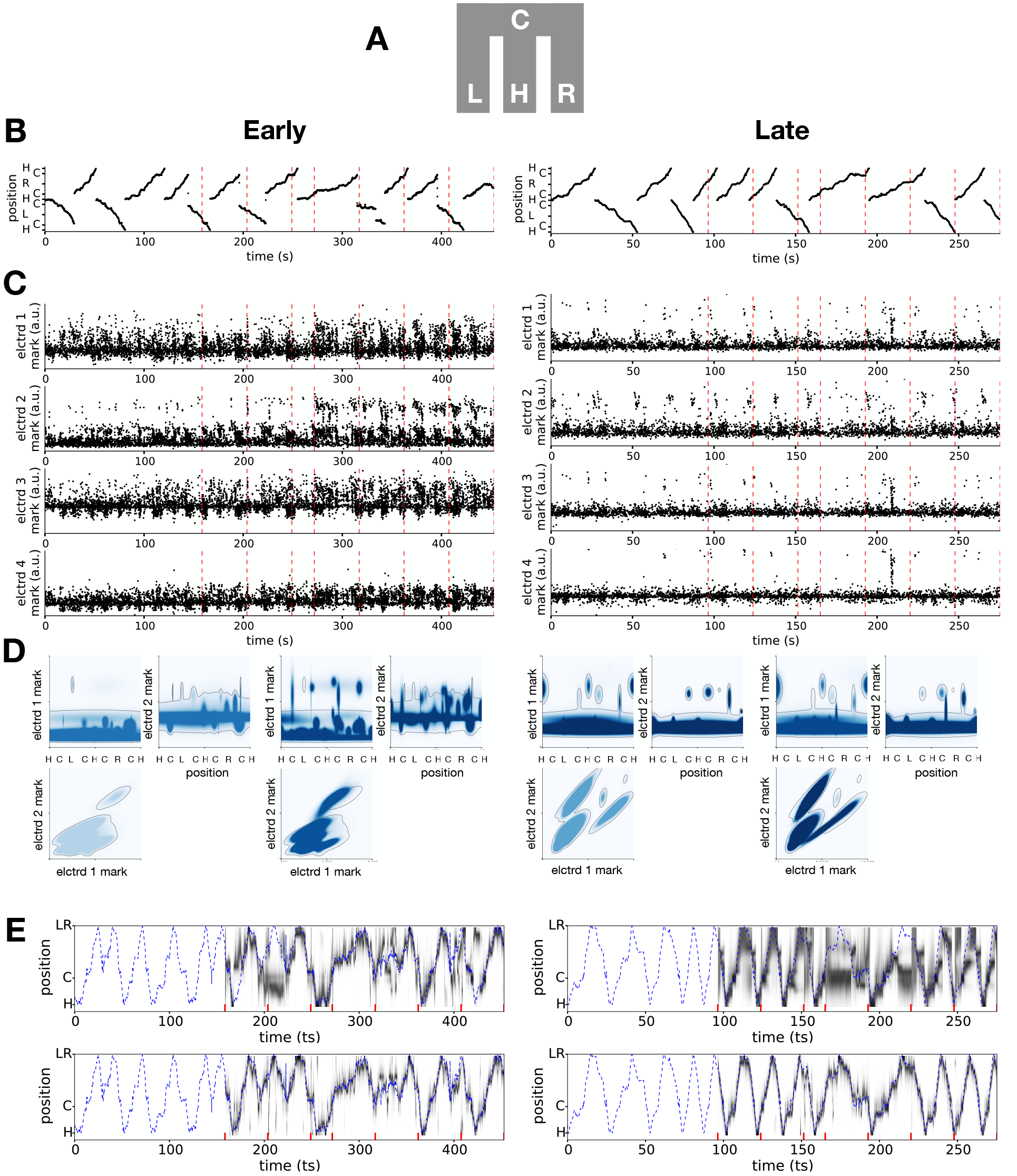
Decoding physical position of rat as it moves through a maze. A) Maze schematic, H: home well, C: choice point, L,R: reward wells on left and right. B) Rat trajectory in a W-maze, displayed in linearized 1-dimensional coordinates. Left and right show early and late exposure to the maze, respectively. Dashed red lines demarcate epochs used for sequential encoding. C) Marks recorded on each of the 4 channels of 1 example tetrode (out of 15) as a function of time. D) The marginalized mark intensity functions at epoch 1 and 5, shown using electrodes 1 and 2 and position. E) Decode example using sequentially updated encoding model. Top row is decoding result using the single tetrode shown above, and bottom row is decode combining all tetrodes. Decoded path displayed by collapsing left and right arms and heading direction to obtain linear displacement from home well.

In the early exposure, our method tracked newly appearing place fields and possibly new neurons. The raster plot in Fig. 8 shows the activity visibly changing around time *t* = 270, and the epoch 1 and 4 JMIFs shown in Fig. 8D shows JOYFULMoGs capturing the newly appeared clusters at later epoch. On a later date, the data from the tetrodes look relatively stationary in both waveform marks and place field structure throughout the experiment, Fig. 8C right.

Example decoded trajectories are shown in Fig. 8E for the early and late exposures. In each day, an example decode using only 1 of the 15 tetrodes is shown above the decode when all 15 tetrodes are combined. Not surprisingly, combining tetrodes improves decode performance. We note that decodes shown are obtained by folding the map and the obtained densities, collapsing the the left and right distinction and the movement direction, so that the coordinates are now just H-C-LR, and represent only the linear displacement from home well. We do this because in the discontinuous scheme we used to build the encoding model, decode metrics like RMSE make little sense.

Fig. 9 shows a comparison between decodes in the early and late exposures. In each row of Fig. 9A), a comparison is made between using 1 tetrode versus combining all tetrodes, with a summary of the spread for 15 individual single tetrode decodes being represented by the error bars. We can see that in the early exposures, decode performance of the single tetrodes increases as more data is accumulated, while in the late exposure, performance remains relatively constant. This effect disappears when combining the tetrodes in this example, suggesting that perhaps only a few tetrodes are becoming more informative later on in the experiment. Fig. 9B) shows a comparison between using an up-to-date SQUP (age of EM=1) versus using a stale SQUP (age of EM=7). For the early exposure, there is a noticeable difference improvement in decoding when the up-to-date decoder is used, while in the late exposure, this difference is less pronounced. These results support the notion that neural representation is less stable earlier in an experience, and that neural representation can also change in relatively short timescales of seconds to minutes.

**Figure 9:**
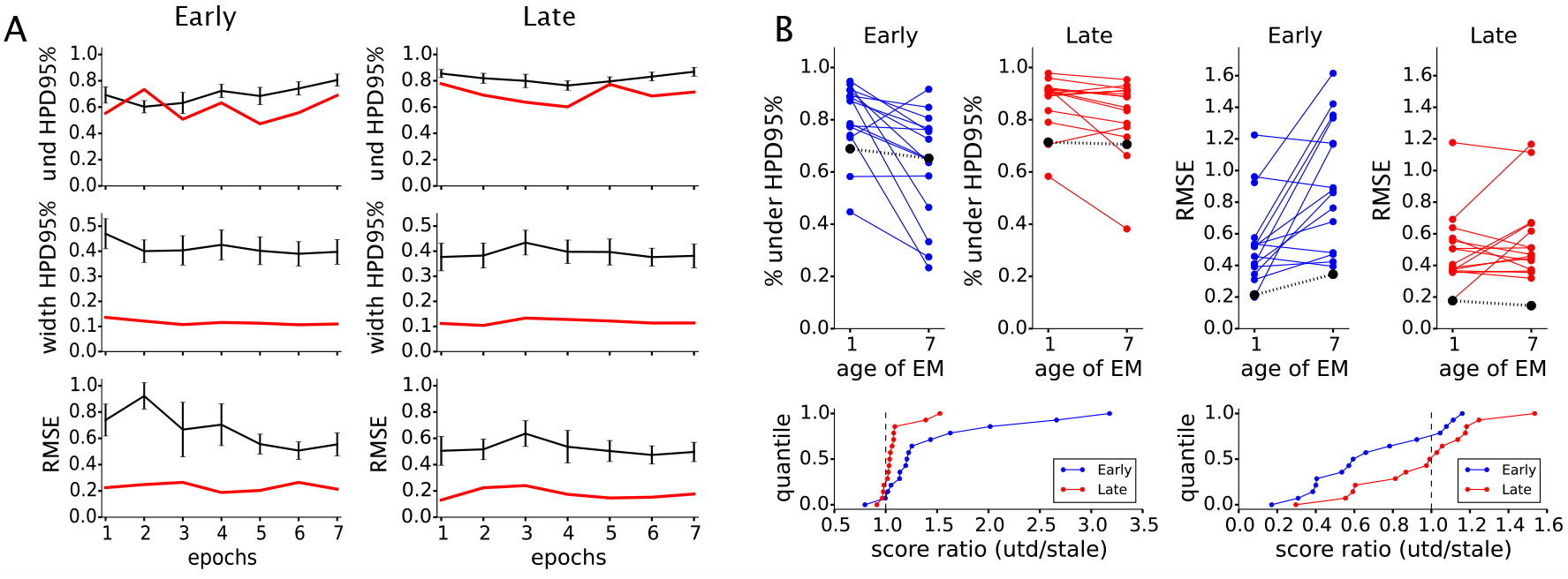
Quality of decode for the early and late exposures. A) From top, time within the 95% HPD region, the width of the HPD region and the RMSE of the physical location from the maximum posterior density for the early and late exposure. 0.1 in RMSE and HPD width corresponds to about 18 centimeters. Black lines are the mean values for the decodes done individually for the 15 tetrodes, with error bars representing 1/4 standard deviation, and the red line the results for decode combining likelihood from 15 tetrodes. B) Comparison of the decoding quality using each of the 15 tetrodes individually of the last epoch, roughly between 405-450 seconds in early exposure and 248-270 seconds in late exposure, using SQUP (age of EM=1), and using the encoding model obtained using data from only epoch 1 (age of EM=7), which is roughly the first 150 seconds in early exposure, and the first 100 seconds in late exposure. The black line is decoding quality using 15 tetrodes combined. C) ratios of the two sets of scores shown in B).

## Discussion

We presented a Bayesian framework for a population model of neural spiking, JOYFULMoGs, that models the joint firing intensity dependent on waveform features and position, bypassing the need for spike sorting and fitting of place fields for each sorted neuron. JOYFULMoGs allows sequential updates to the model so it can incorporate the most recently observed data into the previous estimate of the model parameters, which also allows the model to track changes in the spatial and mark structure of the data, including appearances and disappearances of neurons and place fields. We demonstrated the method on simulated experimental data, showing increasingly accurate parameter estimation in later encoding epochs whether the data is stationary or dynamically changing. We also analyzed recordings from tetrodes placed in the CA3 region of hippocampus of a rat traversing a W-shaped environment, performing a continuous alternation task, and compared performance of early and late periods in training. We observed clear evidence of rapidly changing neural representation in early periods, and our model was clearly able to track and keep the population encoding model up-to-date. In addition, recent work on goodness of fit tests for marked point process models has shown JOYFULMoGs to produce models of activity that fit the data well [Long et al, accepted]. There are, however several outstanding issues that we need to address in order for JOYFULMoGs to be useful in a real-time experimental setting.

First, JOYFULMoGs is still not real-time for most datasets. The most computationally intensive part of our method is the stochastic allocation of spikes into clusters. For data with a relatively small number of spikes, estimation of JOYFULMoGs parameters can be done in real-or nearly real-time on a laptop. However, with larger number of spikes and putative clusters, the calculation of Mahalanobis distances between all spikes and clusters becomes a major bottleneck. This calculation involves *M* × *N* calculations of a distance, which can easily be broken into *M* independent calculations. After all the distances are calculated and spikes assigned to clusters, cluster parameter estimation can all proceed independently. A GPU implementation of Gibbs sampling of Gaussian mixture models has demonstrated [40] over an order of magnitude increase in performance over standard CPU implementations as we have done, suggesting that JOYFULMoGs can be similarly accelerated. Another approach to increasing performance, might be moving to an approximate Bayesian approach like the variational Bayes method to estimate the posterior distributions [41]. This may also allow a more optimal number of clusters to be inferred, or allow us to forgo our batch mode approach to estimation in favor continuous updates may prove more efficient [28, 29].

Second, complicated maze geometries are difficult to represent in 1-dimension, as JOYFULMoGs as presented requires. We presented the results of decoding on a W-maze using a simple linearization scheme, which discontinuously mapped 2-dimensional coordinates onto a 1-dimensional line. This approach may not be satisfactory for more complicated mazes with more than 1 choice point. Further, mazes that combine open areas with narrow maze segments cannot be meaningfully represented in 1-dimension. JOYFULMoGs is easily extensible to accommodate 2-dimensional spatial representations without a substantial increase in computing cost. Decoding using the non-linearized dimensional coordinates of the W-maze have been demonstrated using JOYFULMoGs[in preparation]. There, we assume the spatial covariances of clusters are diagonal so the *x* and *y* components decouple, allowing fitting of the directions one at a time. Because the maze leaves many spatial locations unexplored, it is possible that place fields extend outside the accessible maze locations. However, without observation of such locations, conclusions cannot be drawn about the full extent of the place fields. Therefore, we only concern ourselves with the visible portion of place fields by including dummy trajectory data outside the visible maze, but without any spiking activity. This limits the Gibbs sampler from sampling cluster parameters that would make place fields extend far outside the maze. These simple modifications should extend to more complicated mazes, and mazes that combine narrow segments with wide open areas, allowing real-time analysis of more realistic environments. Future work will explore the use of JOYFULMoGs in such scenarios.

## Acknowledgments

This work was supported by grants from the NIH (MH105174, NS094288) and the Simons Foundation (542971). We also thank Xinyi Deng, Ali Yousefi and Long Tao for helpful discussions.

## Footnotes

1 A notable exception occurs when a cluster’s 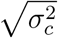 is wider than the width of the sampled trajectory, *W*. 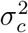 strongly constrains the conditional posterior for λ_*c*_, yet for widths much larger than *W*, the shape of the cluster in that observable space is barely affected by any particular choice of value of 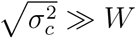 inducing a strong correlation between the samples of 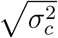 and *lambda*_0,*c*_. However, most informative clusters have 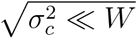 than the whole of sampled space, and this correlation with the intensity is not observed.

